# Towards Computing Attributions for Dimensionality Reduction Techniques

**DOI:** 10.1101/2023.05.12.540592

**Authors:** Matthew Scicluna, Jean-Christophe Grenier, Raphaël Poujol, Sébastien Lemieux, Julie G Hussin

**Affiliations:** Montreal Heart Institute, Research Center, Canada; Département de biochimie et medecine moleculaire, Université de Montréal, Canada; Mila – Quebec AI institute, Montreal, Canada; Département de Medecine, Université de Montréal, Canada

**Keywords:** Visualization, Data Analysis, Bioinformatics, Feature Selection, Machine Learning

## Abstract

We describe the problem of computing local feature attributions for dimensionality reduction methods. We use one such method that is well established within the context of supervised classification – using the gradients of target outputs with respect to the inputs – on the popular dimensionality reduction technique t-SNE, widely used in analyses of biological data. We provide an efficient implementation for the gradient computation for this dimensionality reduction technique. We show that our explanations identify significant features using novel validation methodology; using synthetic datasets and the popular MNIST benchmark dataset. We then demonstrate the practical utility of our algorithm by showing that it can produce explanations that agree with domain knowledge on a SARS-CoV-2 sequence dataset. Throughout, we provide a road map so that similar explanation methods could be applied to other dimensionality reduction techniques to rigorously analyze biological datasets.

## Introduction

Dimensionality reduction techniques such as t-distributed Stochastic Neighbor Embedding (t-SNE) [30], Uniform Manifold Approximation and Projection (UMAP) [14] and Potential of Heat-diffusion for Affinity-based Trajectory Embedding (PHATE) [16], have become widespread tools in the data analyst’s toolbox, achieving popularity in the Machine Learning community and particularly in Bioinformatics. Such techniques can identify structure in high dimensional data by projecting it onto a lower dimensional manifold. When the manifold is 2 or 3 dimensions, the structure can be easily interrogated using ordinary scatterplots. While these methods have informed many data analysis projects, they suffer from an overlooked limitation: there is no obvious way to attribute a datapoints’ embedding to its corresponding input features. Currently, practitioners rely on checking for enrichment of features within groups of points of interest. This is often ad-hoc, and can potentially miss significant features due to cognitive tendencies such as confirmation bias.

We propose a method that can produce such attributions for the t-SNE algorithm. Our methodology is conceptually simple, being based on the well established practice of using model gradients to compute feature attributions [23]. Our algorithm can be added to any implementation of t-SNE, with comparable complexity to the original t-SNE fitting procedure.

In the next section, we describe interpretability methods in more detail, contextualizing ours. We then introduce the constituent parts of our framework: the gradients attribution method and the t-SNE dimensionality reduction algorithm. Then we propose our method to apply gradients computation to the t-SNE algorithm. Then we describe the methods for validating our attributions, describing the results we get when applying our validation methods on the MNIST dataset. Finally, we utilize our attributions to analyze a SARS-CoV-2 dataset – a case study that represents a realistic bioinformatic application. We also performed an additional attribution experiment on the twenty newsgroups dataset that can be found in appendix F. In summary, this work makes the following contributions:

1. Derives the equations to compute the gradient of t-SNE embeddings with respect to each input
2. Produces an algorithm which returns these gradients and is compatible with the Barnes-Hut t-SNE approximation
3. Introduces a novel metric to evaluate dimensionality reduction attribution performance
4. Demonstrates empirical evidence for the methodology on MNIST and SARS-CoV-2 datasets

We have created a Python package that can be installed using the following command:

~~~
pip install interpretable_tsne
~~~

## Background

Previous literature suggests that interpretability is not a monolithic concept, but in fact reflects several distinct ideas [12]. For the purposes of contextualizing this work, we define interpretability as the ability to extract human-understandable insights from Machine Learning (ML) models^1^. One way to ensure interpretability is to use a model which admits a simple explanation by design. Within the supervised learning framework, algorithms have been designed to produce models which are simple enough to be interpretable. These range from classic algorithms like Decision Trees and Sparse Lasso regression [27] to interpretable versions of modern deep learning architectures like BagNets [4]. The limitation of these approaches is that the increased interpretability comes at the expense of model performance.

Practitioners can instead apply post-hoc interpretability methods: which we define as methods that produce explanations of model behaviour after training. There exists many such methods, which can be separated by the kind of explanation they provide: some are *local*, providing explanations specific to each datapoint (e.g. LIME [22], Vanilla Gradients [23]), while others produce *global* explanations of a models activity [26, 19]. These methods can be further grouped based on whether they produce feature attributions, which we define as a score for each input feature which represents the features relative influence on the models behaviour.

Many post-hoc, local, feature attribution methods have been proposed. We can divide these into perturbation and gradient-based approaches. Perturbation-based approaches like LIME [22] and SHAP [13] change parts of the input and observe the impact on the output of the model. The downside of such methods are that they are computationally infeasible when model inference is slow since they require many model evaluations. Gradient based approaches use the gradient (or a modification), to compute feature attributions (e.g. Layerwise Relevance Propagation [2] and DeConvNet [32]). These techniques tend to be much more computationally efficient, but can be insensitive to either data or model [1].

### Gradient Attributions

Within the context of supervised learning of neural networks on classification tasks, techniques have been developed for computing (local) feature attributions. Let *S*_*c*_(*x*) ∈ ℝ be the score function of class *c* given by our classification model, when *x* ∈ ℝ^*d*^ is the input data. Feature attribution methods assign a value to each feature 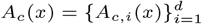. *A*_*c,i*_(*x*) represents how much feature *i* of *x* contributed to the model’s prediction of class *c*.

For this work, we use an attribution method commonly referred to as the vanilla gradient [23]. For our purposes:

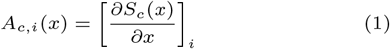

The argument for using the gradients as attribution values provided in [23] is that the above gradients correspond to the weights of the first order Taylor approximation of *S*_*c*_ at *x*. These weights would have a direct correspondence to attributions since the approximation is linear^2^.

In practice, many gradient based attribution methods have been proposed and validated including Integrated Gradients [25] DeconvNets [32] and “Guided Backpropagation” [24]. While such techniques have well known limitations [1, 8], they nonetheless continue to be used all throughout the interpretable machine learning literature.

### The t-SNE Algorithm

The t-SNE algorithm is among the oldest and most influential dimensionality reduction techniques still in widespread use. We provide a sketch of the t-SNE algorithm here. For a detailed discussion of the t-SNE paper, we refer the reader to the original paper [30].

Suppose we have input data *x*_1_, …, *x*_*n*_ ∈ ℝ^*d*^. Denote 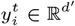 as the embedding for *x*_*i*_ to be produced by the t-SNE algorithm at step *t* in the embedding space with dimension *d*^*′*^ (usually 2 or 3). The t-SNE algorithm updates the 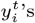 to minimize the Kullback-Leibler (KL) divergence, a measure of the difference between the probability distributions *p*_*ij*_ := *p*(*x*_*i*_, *x*_*j*_) and 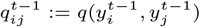:

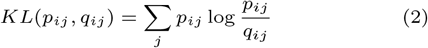

Note that this represents distances between pairs of points in input and embedded space respectively. The intuition is that we want the embeddings in the low dimensional space to recapitulate the distances between points in the high dimensional space. Ignoring optimization hyper-parameters, our embeddings are updated using the following equations:

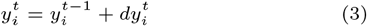

Where:

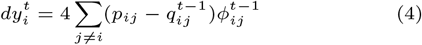

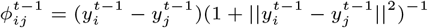

In t-SNE, we update the embedding of each datapoint using (4) until convergence.

Algorithm

The reasoning behind the use of the gradient as a feature attribution method can be used if we consider our score function *S*_*c*_(*x*) to be the output of a dimensionality reduction technique (for embedding dimension *c*) rather than the score of class *c* of a parametric classifier.

Furthermore, the t-SNE update formula (4) is the gradient of an objective function (eq 2) with respect to embeddings *y*_1_, *…, y*_*n*_, and so each *y*_*i*_ is essentially receiving a Stochastic Gradient Descent (SGD) update. We propose inspecting the gradients of t-SNE in the same manner as one would look at gradients with respect to their inputs in relation to supervised classifiers trained also via SGD.

### Computing t-SNE Attributions

In the supervised classification context, computing the gradient with respect to the input is usually very simple, but doing so for t-SNE is more involved, since the relationship between inputs *x*_1_, *…, x*_*n*_ and outputs *y*_1_, *…, y*_*n*_ is less clear. In the following section, we will derive the gradient of each component of a t-SNE embedded point with respect to its input:

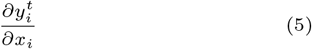

We do not use this gradient directly since we would end up with a set of feature attributions per t-SNE component. This is undesirable since (i) we want only one set of attributions and (ii) the t-SNE components themselves do not have any clear meaning. Instead, we return 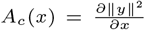. We found that this modification produced attributions that had an easy interpretation: they inform us of how the features of *x*_*i*_ contributed to the overall placement of *y*_*i*_.

### Computing the Gradient of the t-SNE Algorithm

Hereafter, we discuss applying the gradient attribution method to the t-SNE algorithm. We chose this algorithm since it is fairly easy to implement and analyze, and has become widely used within both of the machine learning and bioinformatics communities. We emphasize that our technique could be extended to other dimensionality reduction techniques, provided that they consist of no non-differentiable operations.

We can compute (5) since each of the steps of the t-SNE algorithm are differentiable (we assume that the Euclidean distance is used in the computation of *p*_*ij*_). If we assume that 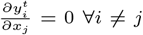, we can compute (5) efficiently using dynamic programming:

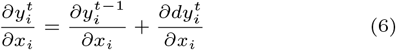

Where:

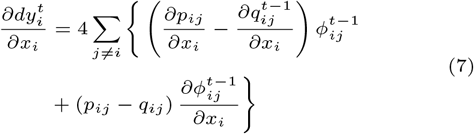

At step *t*, we store 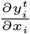 so it can be accessed at step *t* + 1.

This allows us to compute the following:

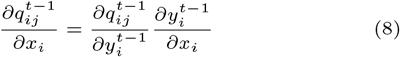

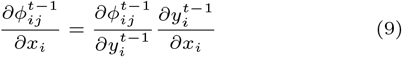

We note that this can be implemented within any implementation of the standard t-SNE algorithm by the addition of a few lines of code. We provide pseudo-code in algorithm 1. See appendix A for the formulas for 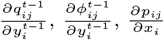, and for the full derivations.

#### Algorithm 1 Gradients for t-SNE

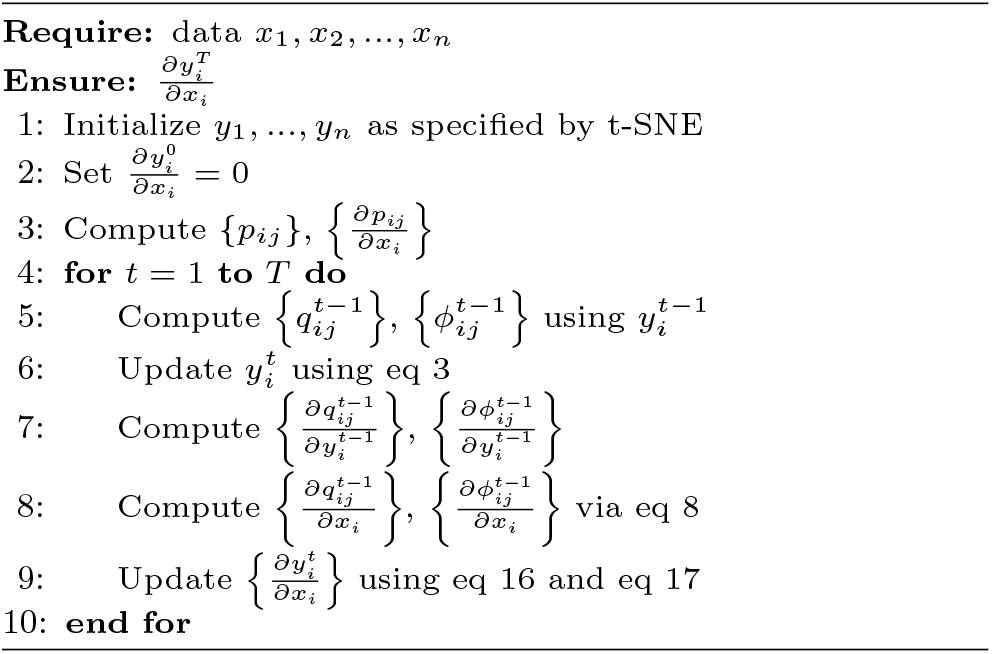

### Barnes-Hut Approximation

Most implementations of the t-SNE algorithm use the Barnes-Hut approximation to speed up computation time from *O*(*n*^2^) to *O*(*n* log *n*) [29]. We show in appendix B how to derive gradients using the Barnes-Hut variant of t-SNE. We note that all experiments reported in this paper were done using gradients of the Barnes-Hut approximation of t-SNE.

## Methods

It is generally very difficult to assess the validity of feature attribution methods, even in their usual supervised classification context [12]. In order to determine whether our attributions were identifying significant features, we performed a series of experiments on synthetic data as well as on the MNIST benchmark dataset. To show real-world applicability of our method, we used our method to identify the mutations driving SARS-CoV-2 evolution using publicly available sequence data. Please refer to appendix G for details regarding t-SNE hyperparameters, attribution processing and performance on benchmarking experiments.

### Simulated Data Experiments

We generated several datasets such that they would have a hierarchical cluster structure whose structure was attributed to a small subset of features. For each data point of a cluster, we translated a small subset of features by a fixed amount. Each cluster was designed such that a small subset of features was translated by a given amount. This set of features differed per cluster, and one cluster did not have any translated feature. We fixed the cluster structure and ground truth feature dependencies, but varied the amount of feature translation that defined the clusters. The details of the data generating procedure can be found in appendix C. After fitting our t-SNE and computing attributions for each synthetic dataset, we took the absolute value of the average of the attributions of all the points in each cluster, and found that, for each simulated dataset, these class-averaged attributions were significantly higher for the ground truth features versus the rest. This was observed even for the cluster that contained no translated features. See appendix C for details of the results.

### MNIST Validation Experiments

We performed a series of experiments using the MNIST dataset. The main idea was to corrupt features based on their attribution values, and then compute the t-SNE embeddings of this corrupted data. If the attributions had detected significant features, then the t-SNE of the corrupted data should be significantly different then the t-SNE fit on the uncorrupted data. We used three separate metrics to quantify the extent of t-SNE structure degradation caused by the data corruption, adapted from metrics used to measure t-SNE quality [10, 11]. These metrics are:

1. <u>Spearman Correlation</u>. The correlation between distances of pairs of embedded points before and after feature corruption. This is a measure of the change of global structure.
2. <u>Adjusted Rand Index (ARI)</u>. We computed the adjusted rand index between clusters generated using K-means clustering (*K* = 10) before and after corruption. This is a measure of the change of cluster structure.
3. <u>10 Nearest-Neighbor Preservation</u>. The average of the 10 nearest neighbors retained by each point before and after corruption. This is a measure of the change of local structure.

We performed our validation experiments on a random subset of 10, 000 MNIST digits. For each experiment, we computed 10 different t-SNEs (random seeds). We varied the percentage of features corrupted from 2% to 18% (in increments of 2%) and report the average over the percentage corrupted.

#### Local, Class-Level, and Global Attribution Validation

On MNIST, we noticed that individual attributions highlighted idiosyncrasies of each digit (see figure 1 A). We noticed that these attributions could be aggregated on a class level, and these saliency maps appeared to be visually meaningful (see figure 1 B). This led us to investigate the validity of these attributions on three distinct levels:

1. *Local* : Attributions produced for each individual digit
2. *Class*: Attributions for each digit class
3. *Global* : Attributions of each feature across all digits

**Fig. 1.**
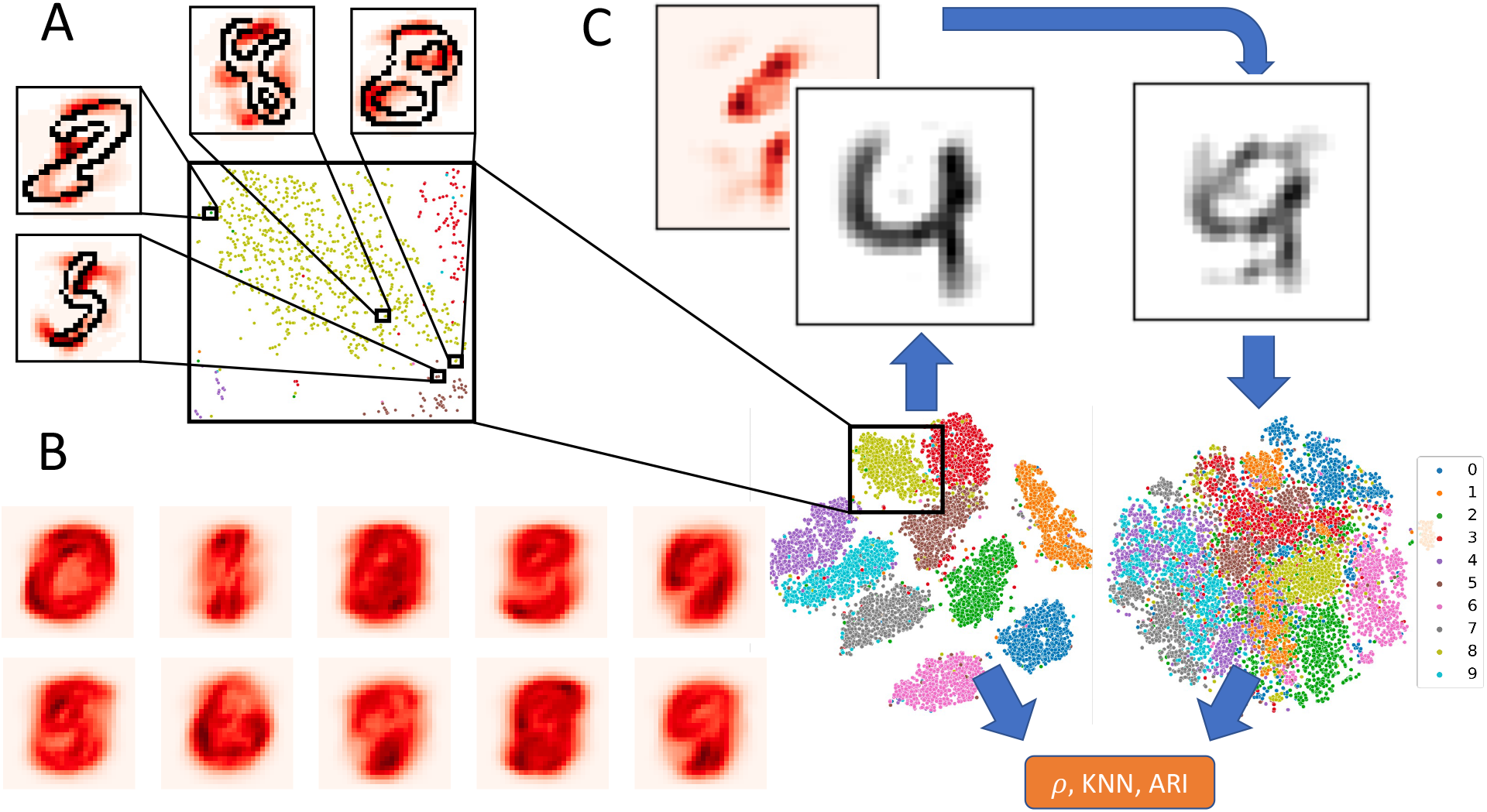
Overall Description of Method and Schematic of Validation Experiment on MNIST dataset. A: We display local attributions superimposed onto t-SNE embedded digits. B: Attributions aggregated (via averaging) within each class. C: We computed t-SNE embeddings and their corresponding attributions using the PCA transformed MNIST digits (left t-SNE plot). We then corrupted the digits based on their attributions (the heatmap and the digit 4 before and after corruption). Note that both digits are projected from PC space using the inverse of the PCA transformation. We then recomputed the t-SNE using this corrupted data as input (right t-SNE plot). We computed metrics such as the Spearman correlation (*ρ*) of t-SNE embedded distances before and after the feature corruption. For A,B and C we projected the attributions from PC space into pixel space by multiplying them by their corresponding PC loadings.

#### Selecting Features to Corrupt using the Attributions

On the local level, we corrupted *k*% of features by corrupting the features within the top *k* percentile of attribution values (in absolute value). On the global level, we corrupted the features that appeared in the *k* percentile of attribution values most often. On the class level, we did the same but for all the points in each digit class separately. Note that on the local level, each digit had a different set of features to be corrupted. On the global level, the same features were corrupted for all digits. On the class-based level, each class had its own set of features to be corrupted, and every digit within a class had the same features corrupted.

Taking inspiration from previous work in the local feature attribution literature [24], we experimented with corrupting features based on attributions produced by positive gradients only and by multiplying the gradients by the inputs (and taking absolute value).

#### Methods of Feature Corruption

For our local and class level attributions, we corrupted each feature by setting all the values to be corrupted by the mean of those values. For the global level attribution validation, we corrupted each feature by removing it from the dataset entirely. To ensure that our results were not biased by our corruption method, we experimented with an additional method of corruption: randomly permuting the values to be corrupted. We replicated all experiments using this permutation corruption method and present the results in appendix D.

#### Baselines

For each level of analysis and each percentage of features to be corrupted, we randomly sampled 10 subsets of features to be corrupted, and computed the change in correlation/10-KNN preservation/ARI to be used as our random baseline. For the individual level attributions, we corrupted a different random subset of features per sample. For class level validations, we corrupted a different random subset per class. For the global validations, we corrupted the same random subset of features for all samples.

For the global level validation, we compared our method to the Laplace Score, a popular unsupervised feature importance [7] method. We computed the Laplace score with respect to both *P* (matrix of *p*_*ij*_ ‘s) and *Q* (matrix of *q*_*ij*_ ‘s) used by t-SNE. In addition, we compared the method to the Fischer Score, which can be seen as the supervised version of the Laplace Score [7]. We also compared the method to the top principal components, representing a variance-based control. For the class level validation, we computed a “class-based” Laplace Score by re-computing the *P* and *Q* matrices on each class subset and then computing the Laplace Scores. To compute the Fisher and Laplace scores, we used the python package `scikit-feature`.

At all levels, our final baseline was to select features using the absolute values of those features. For the class-based and global experiments, we selected features in an analogous manner as was done with our attribution based methods, except that we substituted the feature values in place of the attributions. Refer to figure 1 C for a schematic of the validation experiment.

### SARS-CoV-2 Case Study

In order to demonstrate the practical utility of our method, we used it to investigate SARS-CoV-2 sequence data. The project has ethical approval from the Ethics Board of the Montreal Heart Institute, Project 2021-2868. We downloaded a globally representative sampling of 3, 064 SARS-CoV-2 via Nextstrain [6] accessed January 26 2023. The sampling was done between Dec 2019 and Jan 2023. We intersect these with the condon-based alignement of GISAID [5] from march 15 2023 resulting in a final dataset of size 2, 374 (EPI SET ID EPI SET 230418kp). The down sampling is due to the filtering perform by GISAID on missing data during the alignement process. We then recode as missing data any deletion greater than 12 nucleotides. We note that our dataset may be biased due to the sampling done by NextStrain. We derived the allele states from the Wuhan ancestral sequence (Gisaid ID: EPI ISL 402124). The multiple sequence alignment (MSA) was performed using an optimized multiple sequence alignment procedure made by GISAID using MAFFT [9]. Each observed mutation or deletion at each position was encoded as a 1 if that mutation or deletion was present in the sequence and 0 otherwise. We ignored mutations or deletions that only occurred once in our dataset. Finally, we ignored any mutations occurring in the first or last 100 positions as these are less covered by the sequencing and thus of low quality. This left us with 33250 mutations and 3359 when removing the reference allele.

For each sequence, we obtained Pangolin annotations [20, 18] from GISAID, and used these to classify each sequence as belonging to either “Alpha”, “Beta”, “Delta”, “Gamma”, “Omicron”: BA.1, BA.2, BA.4, BA.5 and BQ as designated by the World Health Organization (WHO). We labelled recombinant lineages such as “XBB” separately.

We downloaded representative genetic markers for each lineage from outbreak.info [28]. We removed markers containing deletions, since we were unable to identify the exact genetic positions of them.

## Results

### Qualitative Results on MNIST dataset

We found that on the local level, our t-SNE attributions highlighted digit idiosyncrasies (see fig 1 A). On the class based level, we found that the digits highlighted pixels that varied within classes, but also seemed to suggest which digit classes would cluster together in the resulting t-SNE. For example, looking at the class averaged attributions in B of figure 1, we see that the averaged attributions of the 4’s look very similar to those of the 7’s and 9’s, and indeed these three clusters appear next to each other in t-SNE space (almost forming their own “super cluster”. We observe the same pattern between the 3’s, 5’s and 8’s.

### Local, Class-Level, and Global Attribution Validation Results

For each of the local, class, and global level, we found that our methods significantly outperformed the random baseline and were on par with or superior to the other baselines.

For the individual level baseline, we experimented with using only positive attributions. We found that these performed worse then just using the attributions themselves, and so we ignored them in subsequent experiments. We found that multiplying the attribution by the absolute feature value yielded the best 10-NN preservation (averaged across corruption %) at 0.20 *±* 0.0024 versus the second-highest value of 0.28 *±* 0.0035. Similarly the ARI was 0.36 *±* 0.0240 versus second best value of 0.38 *±* 0.0129. The Spearman correlation was a close second to the feature value baseline: 0.35 *±* 0.0361 vs 0.32 *±* 0.0510.

For the class level experiments, we found that the attribution alone either outperformed or were on par with all other baselines (Spearman 0.49 *±* 0.0992 vs 0.50 *±* 0.0657, KNN Preservation 0.49 *±* 0.0034 vs 0.50 *±* 0.0019). For the global level experiments, we found that both our gradient attribution-based methods either outperformed or performed on par with the other baselines. The full results can found on figure 2 and on tables S2, S3, S4 in the supplementary materials. We highlight that our method is on par with other well established methods from the feature importance literature, despite being developed from the local feature attribution framework.

**Fig. 2.**
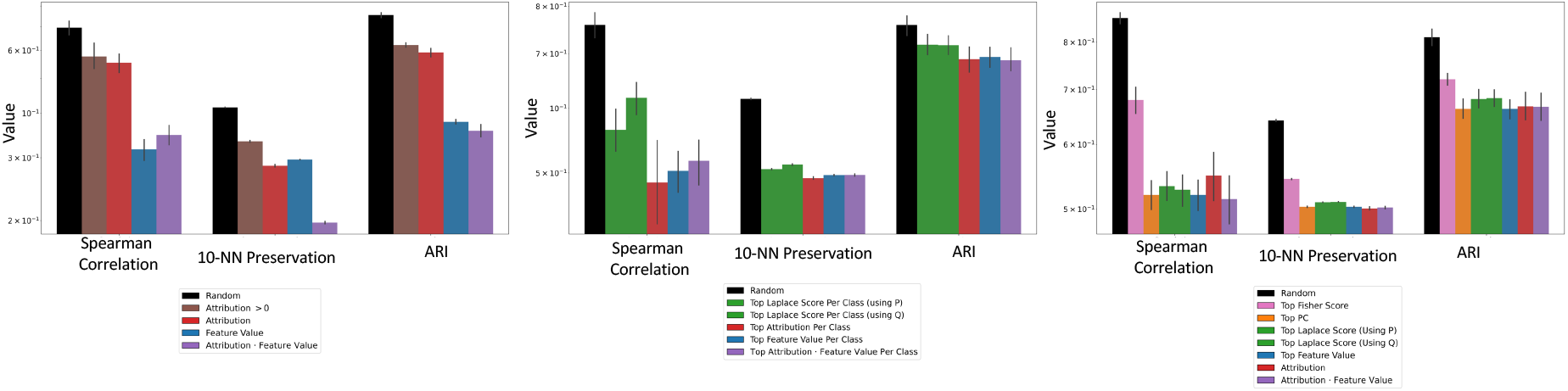
Individual, Class, and Global-level Attribution Validation Experiments performed on MNIST. *(Left=Local)* we corrupted each feature using the mean of the sampled features to be corrupted. At each level, we compared the t-SNE embeddings before and after feature corruption using 3 metrics: the Spearman correlation, 10-nearest neighbor preservation and adjusted Rand index (the y-axis). Note that lower values of each metric means that the corruption affected the embeddings more. Our baselines are (from left to right) random corruption, using only positive attributions, using only the attribution, using only the absolute feature values, and multiplying the attribution by the absolute feature values. *(Middle=Class-based)* we corrupted each feature using the mean of the sampled features to be corrupted. (from left to right) random corruption, using the class-based Laplace Score on matrices *P* and *Q*, using the attribution, using the feature, or multiplying the attribution by the feature. *(Right=Global)* we corrupted each feature by removing the features to be corrupted. Our controls are (from left to right) random corruption, using the Fisher score (supervised feature importance control), using the top principle components (variance-based control), using the Laplace Score on matrices *P* and *Q* (unsupervised feature importance control), using the absolute value of the feature, the attribution, or multiplying the attribution by the absolute feature value. The error bars are 95% bootstrap CIs over the random seeds (and over sampling for our random baselines) computed using seaborn.barplot.

### SARS-CoV-2 Case Study

We wanted to see if our t-SNE attribution method would assign high attribution to the mutations or deletions that we expected to be lineage defining. In order to do this, we needed to ensure that our t-SNE recapitulated the relevant lineage structure. We did this by inspecting a scatterplot of the t-SNE embeddings.

### Using t-SNE Attributions For Quality Control

Our initial SARS-Cov-2 encoding scheme did not yield t-SNE embeddings that clustered based on the WHO designations. This led us to perform an analysis of the t-SNE embeddings using our proposed attribution method. When we compared the attributions averaged within clusters generated by DBSCAN, we found that for several of the clusters, the attribution score was positively correlated with the missingness frequency. Given that the attributions were identifying missing values as the cause of certain clustering patters, we chose to impute this missing data as the reference genotype. For full details of our attribution based QC, see appendix E.

When we computed a t-SNE of our imputed SARS-CoV-2 sequence dataset, we found that the sequences did generally cluster based on their WHO designation. As can be seen in the t-SNE scatterplot of figure 3 A., most clusters correspond to a single lineage, with sub-lineages appearing as nearby sub-clusters. Note that there are some devations in the observed scatterplot. For example, the clusters corresponding to sublineages of BA.5 (BA.5.1 and BA.5.2) do appear on opposite sides of the t-SNE scatterplot.

**Fig. 3.**
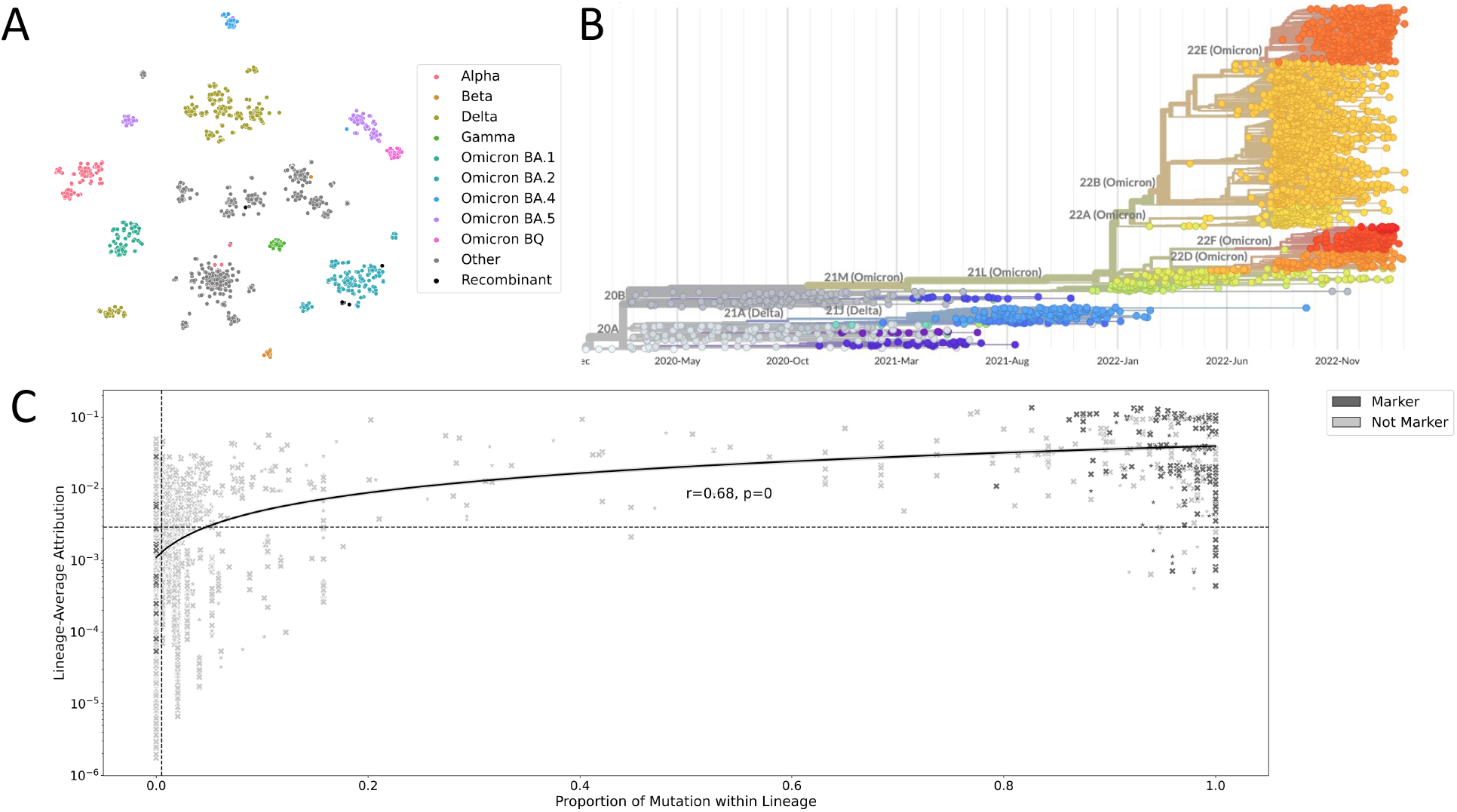
Finding genetic markers for SARS-CoV-2 lineages. A: the t-SNE embeddings of our SARS-CoV-2 dataset. We added the additional “Omicron BQ” and “Recombinant” categories. B: Phylogenetic tree fit via Nextstrain on sequences used in this study [6]. C. We averaged the attributions per mutation for each lineage, and plotted them against the mutation/deletion frequency. We colored the points based on whether they were a marker gene as determined by outbreak.info. Points marked as * are synonymous, and points marked as X are non-synonymous. The dashed lines on the x and y axes indicate the 90th percentile for the mutation/deletion frequency and averaged attribution respectively

### Identifying Genetic Markers from Lineage-Averaged Attributions

Motivated by the apparent utility of class-averaged attributions when used with MNIST, we averaged the attributions of each mutation/deletion per lineage and compared this to the mutation/deletion frequency. Note that the mutation/deletion frequency is a feature average since we encoded each mutation/deletion as a binary variable.

We chose the 90th percentile to be our threshold of significance when identifying mutations/deletions based on attribution scores or mutation/deletion frequency. Of the 267 markers, we found that 251 could be identified by having significantly high mutation frequency, while 229 could be identified by having high attribution. However, 3 markers were identified using attributions that had low frequency. 13 markers could not be identified using either the attributions or mutation frequency. This can be seen on Fig 3 C, where the markers detected by attributions and not feature means appear in the top left quadrant, and the 13 markers not detected by either method appears in the bottom left quadrant.

The attribution based method uniquely identified the Omicron BA.1 marker Spike:G142D and the Alpha and BA.2 marker ORF8:L84S. Both methods missed Spike:N440K (BA.2, BA.4, and BA.5 marker) as well as Spike:N679K (BA.1, BA.2, BA.4 and BA.5) and ORF8:L84S (for Beta, Delta, Gamma, BA.1, BA.4, and BA.5). Of the 25 markers missed by our attribution based method, 21 of them were markers of Gamma, 2 from Beta, and 2 from Omicron BA.4. We suspect that our approach had difficulty identifying these markers because their lineages were the least frequent within our dataset (among the sequences that had markers). In fact, the dataset contained only 34, 49, and 58 sequences of Gamma, Beta and BA.4 respectively.

Finally, we note that our highly attributed mutations were corroborated in the literature. For example, a previous study [17] identified 25 “Haplotype defining” mutations (highly predictive of SARS-CoV-2 evolutionary structure). 24 of these positions were highly attributed by our method.

## Discussion

To the best of our knowledge, this is the first application of a feature attribution method to any dimensionality reduction algorithm. Furthermore, we develop a novel validation method, and provide a biologically relevant demonstration. We note that the algorithm presented provides feature attributions with respect to a given t-SNE embedding. Therefore, any insights yielded by the attribution scores only represent “true signal” from the data insofar as the t-SNE embedding has modeled the data appropriately. This is demonstrated in figure 1 C, where the t-SNE embeddings for the 3 and 5 digit classes are very similar, and indeed the t-SNE embedding has both digit classes adjoined, and not fully resolved on their own. Our method can identify such algorithmic artifacts, which can be useful for practitioners who want to understand why their embeddings appear a certain way, without having to do ad-hoc feature enrichment analysis.

In practice, we suggest that users analyze the attributions of high quality t-SNE embeddings. There exist metrics that quantify t-SNE embedding quality [10, 11]. We suggest that practitioners use them to filter out potentially problematic t-SNE embeddings prior to attribution analysis.

Our MNIST data exists in a human understandable space, and so we can visualize our attributions at each level, and this provides a sanity check for our method. Qualitatively, we found that our attributions yielded human understandable insights about the variation of individual digits and the defining characteristics of each MNIST digit class that were recapitulated by the t-SNE embedding.

On all levels, we found that the attributions produced by our methods significantly outperformed random feature corruption. We are not surprised that our method did not always outperform baselines, particularly at the class-based and global level, given that our method is a local feature attribution method. We hope that the development of this method could inspire future research, to eventually develop less noisy variations of our approach.

We note that throughout this work we implicitly assume that the ground truth feature dependencies are somewhat sparse (i.e. only a few features driving the structures recapitulated by t-SNE). This assumption appears to hold for the datasets used here. In cases where the data exhibits complex relationships between features and structures, it is not clear if one should use feature attribution methods since such relationships may not be well represented by per-sample per-feature scores.

In the SARS-CoV-2 application, we further found that aggregating lineage-averaged feature attribution scores identified significant variations within SARS-CoV-2 lineages. We note that other methods exist for finding markers mutations for SARS-CoV-2 variants, and these have been used extensively to analyze SARS-CoV-2 data in the last 3 years. This is precisely the information that we wanted to leverage to confirm the validity of our attributions in a biological application. In contrast, the ground truth of attributions in other biological modalities such as transcriptomics, metagenomics, or metabolomics can be harder to establish, making the evaluation of attributions trickier. Our approach is not meant as a replacement for other methods, but the ample domain expertise in this field made it appropriate a point of reference to assess our method. Nonetheless, our method could be used on sequence datasets from future waves to identify quickly new sub-lineages arising and to identify outlier sequences to be removed.

We anticipate that this work can be extended in multiple ways. First, we would like to see this method applied to more real-world biological data science applications (including gene expression, protein interaction, metagenomics and metabolomics). We are particularly intrigued by applications in the Single-Cell RNA transcriptomics domain, where t-SNE analysis is particularly popular [10]. However, since ground truth is generally missing in these applications, simulation work will be needed to validate the approach [31]. The algorithmic complexity of our method scales roughly linearly in terms of the number of input features when compared to the usual t-SNE. This is due to additional computations of large, multidimensional arrays. Increasing the efficiency of these computations is a second promising extension. Permutation-based attribution methods such as SHAP [13] have nice mathematical guarantees, but a naive application of such methods would require an infeasible number of model evaluations. Being able to adapt such methods to this problem setting represents a third possible future direction for this research.

## Conclusion

We propose a feature attribution method designed for t-SNE. To the best of our knowledge, this represents the first such attempt for any dimensionality reduction algorithm. In fact, this is also the first attempt to do attribution of a non-parametric machine learning algorithm. We argue that since both methods are optimized via SGD, the gradient with respect to inputs represent the same thing.

We developed a method that evaluates the validity of our approach. Our method quantifies the feature attribution performance by comparing the extent of degradation of t-SNE embeddings post-corruption. We chose baselines from the unsupervised feature importance literature. We also compared our method with feature enrichment baselines, and with appropriate random baselines.

We demonstrated our algorithms correctness using synthetic data, where we knew the significant features available. We then evaluated our algorithm on MNIST. Here, we did not have the significant features known in advance, but were able to provide evidence for our approach using our validation method. Finally, we demonstrate the utility of our method via a SARS-CoV-2 case study, finding that in all cases our approach yielded unique insights that could help a data scientist better understand their t-SNE plot. We hope that this work can serve as the foundation for other works investigating the use of feature attributions for dimensionality reduction algorithms.

## Derivation of t-SNE Gradients

We derive the equations for the gradient of the t-SNE [30] output embedding with respect to the input data. Suppose we have datapoints *x*_1_, …, *x*_*N*_ ∈ ℝ^*D*^. We denote *x*_*i,d*_ as the *d*th feature of the *i*th datapoint. Within the context of supervised classification, the gradient attribution method [23] is defined as follows. For a given score function of class *c*: *S*_*c*_(*x*) ∈ ℝ, we define the attribution for *x*_*i*_ as:

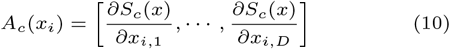

The t-SNE algorithm is a dimensionality reduction technique. Given our data, it will return embeddings of dimension *C < D*. We can think of our output dimension as a score function, so we can write:

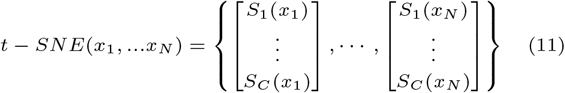

To keep the notation consistent with the original t-SNE paper, we denote *y*_*i,c*_ = *S*_*c*_(*x*_*i*_) and *y*_*i*_ = [*y*_*i*,1_, *… y*_*i,C*_]. The t-SNE function is iterative (over steps 1, *…, T*). We denote the output embedding at step *t* as 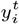. Ignoring the optimization terms, we have that:

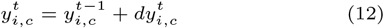

Where:

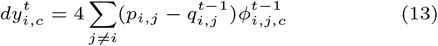

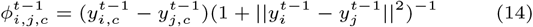

In this set-up, we notice that we could compute:

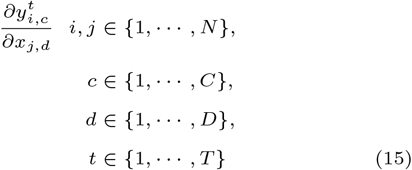

We restrict our interest to **only gradients of an embedding with respect to their corresponding input data point**:

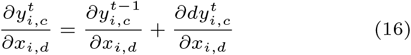

We use the chain rule on 13 to obtain:

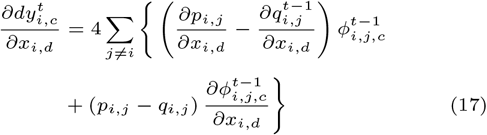

At step *t* − 1, we store 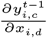 so it can be accessed at step *t*.

This allows us to compute the following:

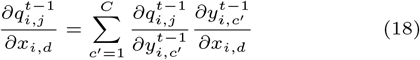

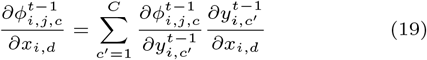

We now derive the gradients for 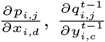 and 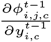.

From the t-SNE paper:

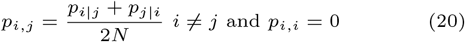

So we must differentiate w.r.t. both components.

If we let 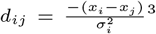 and 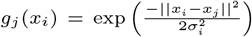, then:

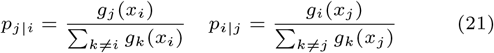

We can apply the quotient rule to differentiate this:

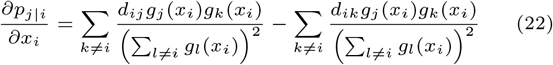

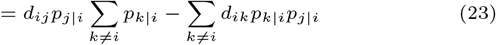

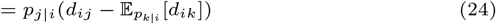

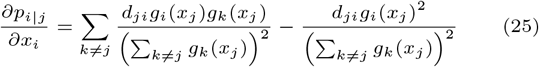

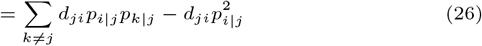

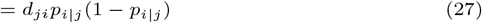

We next derive 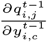

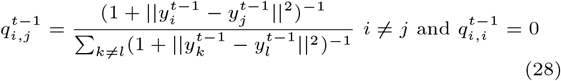

Now let *g*_*j*_ (*y*_*i*_) = (1+||*y*_*i*_ −*y*_*j*_ ||^2^)^−1^ then, using the quotient rule (suppressing unneeded indices):

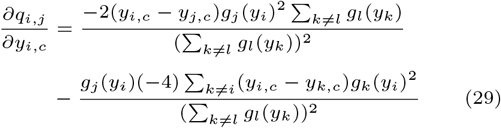

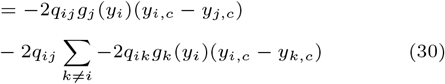

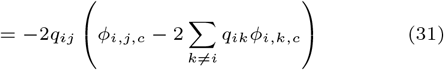

Finally, for *ϕ*_*i,j,c*_

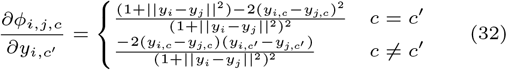

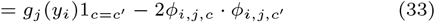

## Derivation of Barnes-Hut Approximated t-SNE Gradient

We show how we can use the Barnes-Hut approximation on the t-SNE gradient (attribution) function. Recall that our attributions can be computed as:

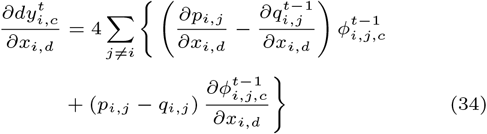

We can rewrite 34 as:

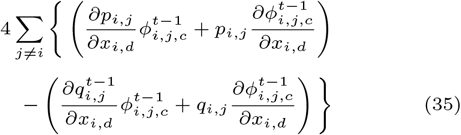

We can break the computation up into 2 parts. For the positive half, we notice that:

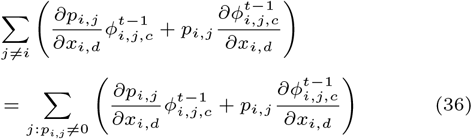

This is because 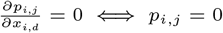. As is done in [29], we can use a sparse *P* matrix to reduce this computation to *O*(*n* log *n*).

For the negative half, we assume that we computed a quad-tree as per [29]. We can approximate this term using the summary embedding per quad-tree cell (denoted as *y*_*cell*_):

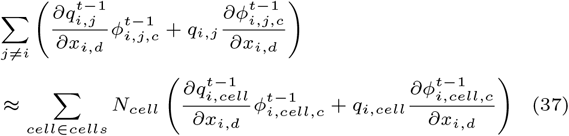

Where:

*N*_*cell*_ = Number of points in the cell

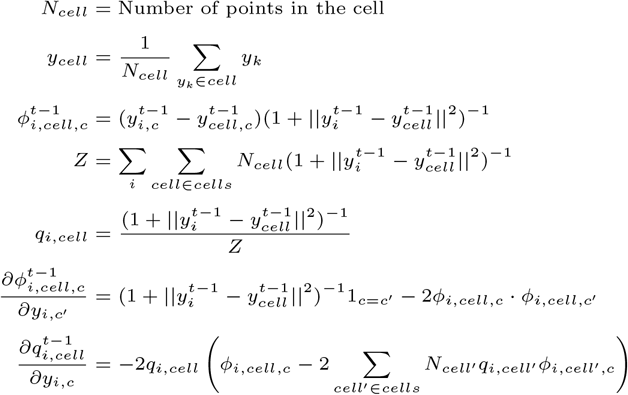

We can improve the efficiency of this calculation by reusing our quad-tree and *Z* term computed during the t-SNE objective calculation.

### Synthetic Data Experiment

We generated 10 dimensional random normal data containing 4 classes. Each class was distinguished by translation of a feature by a fixed amount (enrichment). We refer to this level of enrichment as “Effect 1”. Feature 1 of class 1 and 2 were always enriched relative to class 3 and 4. Feature 2 was enriched by a fixed amount in class 1 relative to class 2, and feature 3 was enriched by the same amount in class 3 relative to class 4. See figure 4 for an illustration of this. We refer to this level of enrichment as “Effect 2”. We varied “Effect 1” from 2,4,6 and “effect 2” from 1,2,3,5. We set “Effect 1” *>* “Effect 2” to make the structure heirarchical. We generated 10 datasets per combination and ran t-SNE with 10 different initializations on each of these. For each dataset, we averaged the attributions over each t-SNE and over all datapoints from the same class and took the absolute value.

**Fig. 4.**
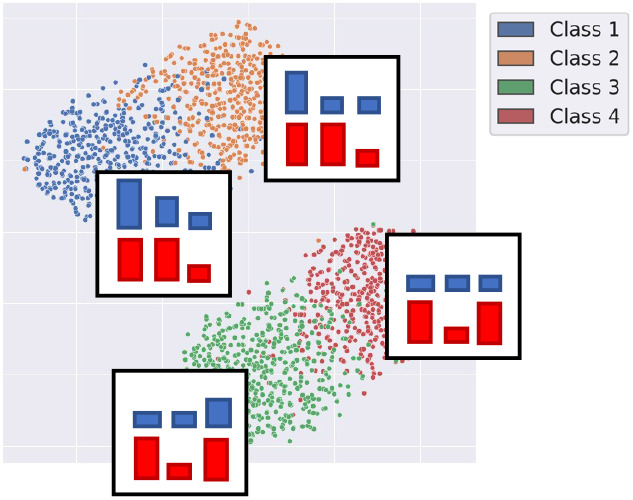
The scatterplot of the t-SNE embeddings fit to synthetic data where “Effect 1” = 6 and “Effect 2” = 3, where each embedded data-point is coloured by its class label. For each class and for the first 3 features, we display the feature value (top, in blue) and absolute value of the expected class-averaged attribution (bottom, in red). Note that we do not include the other 7 features since they are not enriched in any class, and we don’t expect them to have high attribution.

Based on the data generating regime, we would expect the following features to have the highest attributions:

1. feature 1 for all classes
2. feature 2 for class 1 and 2
3. feature 3 for class 3 and 4

We performed a Mann-Whitney U-test to detect if the class-averaged attributions of these significant features were significantly greater then those of the remaining features. We performed our analysis on attributions coming from classes 1, 2, 3 and 4. From table S1 we can see that the class averaged attributions are significantly higher for the known important features versus the rest. We note that this holds even for class 4, which has no feature enrichment.

### Putting this Experiment into Biological Context

The synthetic datasets were chosen to demonstrate that, if a small number of features are causing the observed structure, then our method will (mostly) identify those features. We do not guarantee that this adequately represents all real-world data or covers a wide range of scenarios. While the synthetic datasets contain cluster structure, we also envision synthetic datasets that contain trajectory structure (which is often found in single-cell transcriptomics data). t-SNE tends to have difficulty modeling such data, so remains to be seen how our method would perform in that context.

### Perturbation Corruption Experiments

We repeated the local, class-based, and global level attribution experiments by permuting the values of each feature for all samples whose feature was to be corrupted. The results are presented in tables S5, S6 and S7. Note that the results are very similar to figures 2, except that the improvement from using feature-based removal is more pronounced on the individual level. We believe that this is an artifact of the corruption process, since permuting large values with each-other would have a more detrimental effect on the t-SNE versus permuting smaller values. Ideally, we would like to have “removed” the features, but we could only do so for the global level attribution experiments.

### Using t-SNE Attributions For Quality Control

As mentioned before, our initial SARS-Cov-2 encoding scheme did not yield t-SNE embeddings that clustered based on the WHO designations. See in figure 5 A for a scatterplot of the t-SNE embeddings. That coding scheme included an additional column per position that was set to 1 if the position matched the reference and was set to 0 otherwise. Therefore, we can infer which positions are missing based on if all corresponding columns are 0. This motivated us to investigate the cause of this, providing another realistic use case for our methodology.

**Fig. 5.**
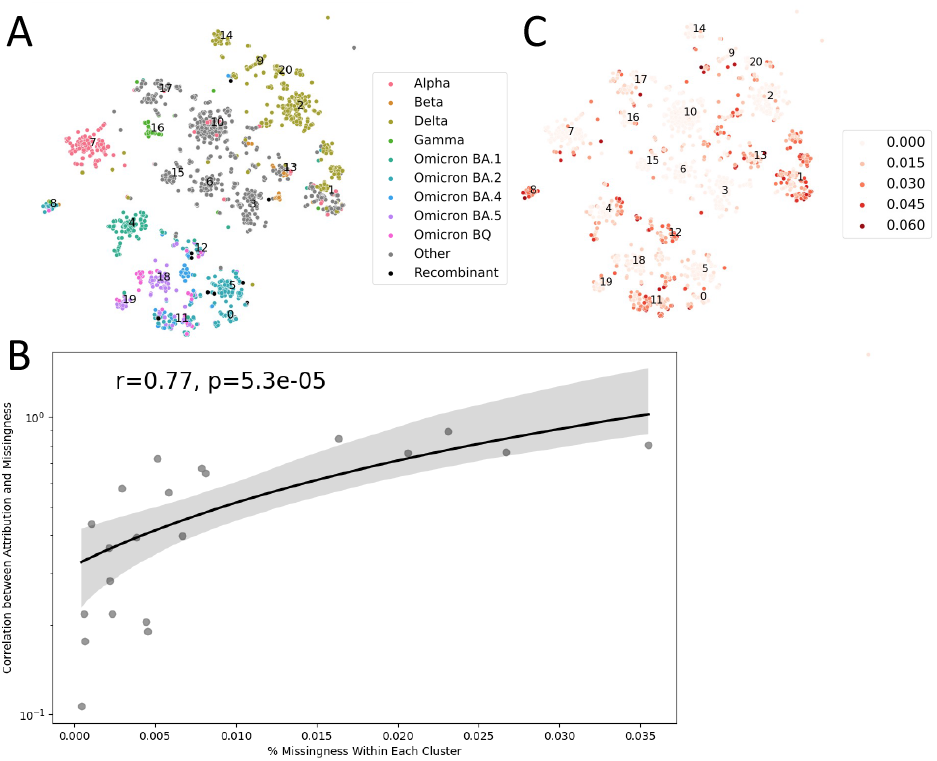
Using Attributions of t-SNE to identify the cause of malformed embeddings. A: the malformed SARS-CoV-2 t-SNE embeddings. The centroid of each DBSCAN cluster is labelled with the cluster number. B. The correlation between the cluster-averaged attribution (y-axis) and the frequency of missing positions per cluster (x-axis). We also plot a line of best fit and display the correlation and p-value. Note that the y-axis is log-scaled. C: The proportion of missingness of each sequence displayed over the t-SNE embedding scatterplot.

We first ran a DBSCAN clustering on those t-SNE embeddings. We then averaged the attributions of all the points within each cluster. We suspected that the cause of the malformed t-SNE embeddings was due to data missingness. We noticed that we could leverage our attributions to provide some evidence for this.

For each cluster, we investigated the relationship between the average attributions at each position and the frequency of missingness for each datapoint. If the missingness did not affect the cluster position, then we would expect a low correlation here. We found that this correlation was high for some clusters and low for others. We suspected that clusters containing the most missing positions would be the most impacted by the missingness (i.e. have the strongest positive correlation between attribution size and missingness frequency). Indeed, when we plot these quantities against each-other for all 20 clusters, we observed exactly this (see figure 5 B). Finally, since we had the missingness data available to us, we plotted the frequency of missingness per sequence and found that several clusters did appear to be highly enriched in missingness. In particular, clusters 1, 8, 11, 12, and 13 contain significantly high amounts of missingness, without a dominant lineage. The correlations of these clusters are 0.89, 0.81, 0.76, 0.76, and 0.85 respectively.

We note that removing the reference columns is equivalent to doing a kind of imputation where we substitute the missing data values for the reference values. When we recomputed the t-SNE after doing imputation we found that the embeddings clustered based on lineage, providing further evidence towards our initial suspicion. This can be seen in figure 3.

While we could have done this analysis using just the missingness information available to us, we emphasize that the attributions still would have correctly identified the positions that are often missing, providing a signal that would be useful here.

### Twenty News Groups Application

To demonstrate that our method works for data coming from very different modalities, we performed an analysis on the twenty newsgroups dataset. This dataset consists of ≈18000 newsgroups posts on 20 topics. For our purposes we further coarse grained out topic categories into the following: “Vehicles”, “Sports”, “Computers”, “Ads”, “Religion”, “Politics”, “Medicine” and “Space”. For each post, we removed all punctuation, “stop words”, and any words containing non-alphabet characters. We used the nltk python package for our data pre-processing [3].

We computed the document vector by taking the average of Word2Vec document embeddings [15] gensim implementation [21]. We performed our analysis on a random subset of 12000 posts, keeping only sentences with at least 25 words and at most 750. We projected our 300 dimensional document embeddings into 50 dimensions using PCA.

We performed t-SNE using the same hyperparameters as was done with the other datasets. Figure 6 A shows that, for the most part, posts from the same topic clustered together. We also see that the attributions highlighted words that were high associated with the post topic. Similarly, when we averaged the attributions per topic, we found that the highest attributed words consisted of jargon specific to the topic, or other words that were similarly highly associated with the topic. In figure 6 B we contrast the highest scoring words with the highest frequency words. Not surprisingly, the highest frequency words consisted of popular, generic words that were largely non-topic-specific.

**Fig. 6.**
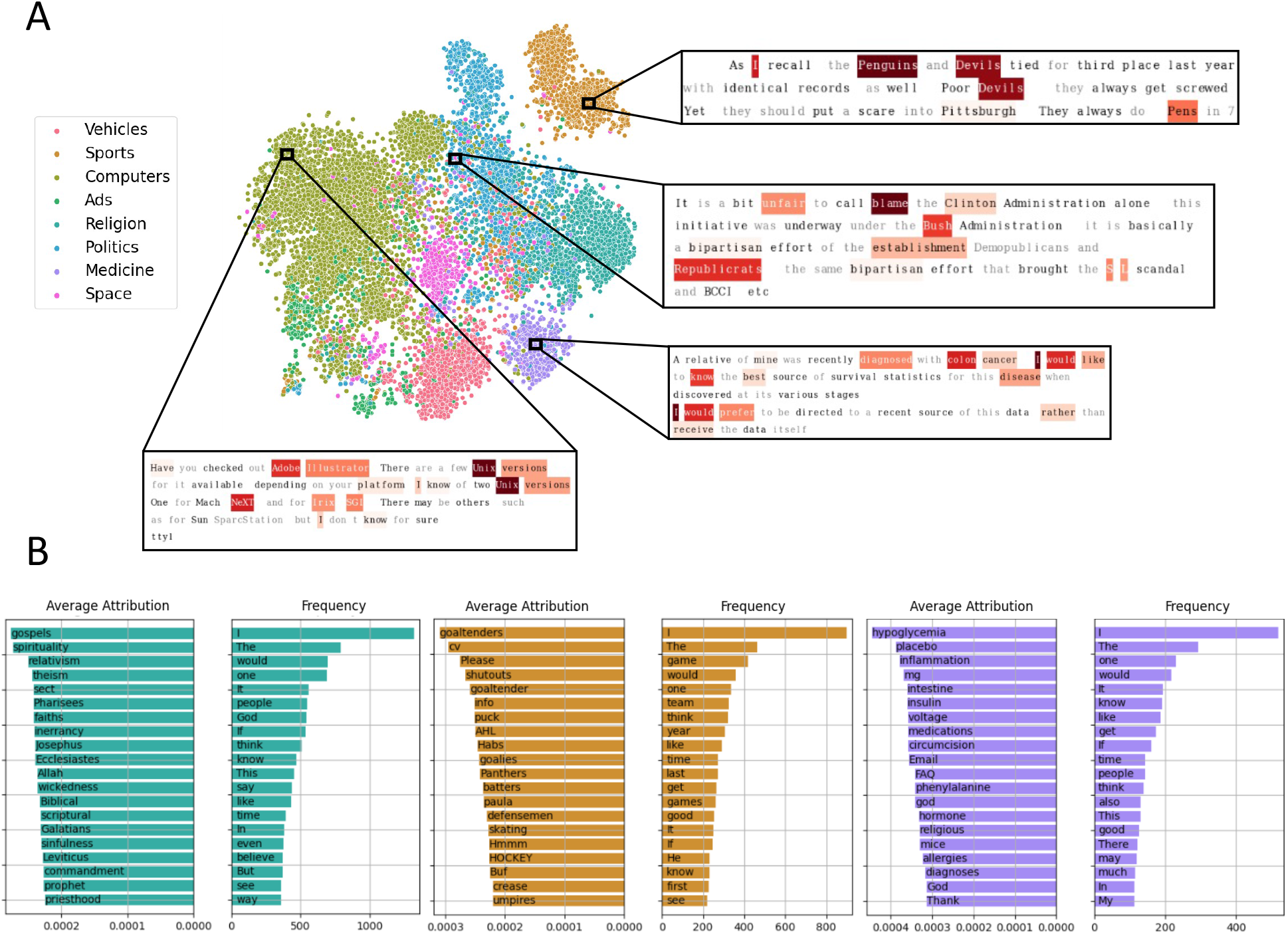
t-SNE attribution analysis performed on the 20 NewsGroups dataset. A. The t-SNE group according to the conversation topic, as expected. When looking at attributions per conversation, we see that the highest attribution words tend to be strongly associated with the conversation topics. B. The average attributions per words (left side of bar plot) compared with the highest frequency words (right side), labelled with the word. The bidirectional bar plots use the same topic color mapping as the t-SNE.

### Gradient Computation Details

We performed our t-SNE experiments using the following hyper-parameters:

1. perplexity = 30
2. number of iterations = 1000
3. the number of iterations of early exaggeration = 250
4. early exaggeration = 4
5. learning rate = 500

### We did not experiment with other t-SNE hyper-parameter settings

We processed the attributions in the following way: first we extracted the attributions computed at the 250th step. We then removed any NaNs and clipped these so they would be in the range of -1 and 1. Finally, to convert the attributions with respect to the PC variables to those of the original inputs, we multiplied them by the PCA loadings matrix. When reporting attribution values or averages, we report the absolute value of the attribution(s).

### Numerical Instabilities

We note that our proposed attribution computations are generally numerically stable. However, a small number of gradients may be very large in magnitude (and possibly represented as NaNs). For example, for one of our MNIST runs, 0.36% of attributions were *>* 1 (0.19% *>* 10) and 0.36% were NaN. For our SARS-CoV-2 data, 0.042% of attributions were NaNs and 1.07% of the data was *>* 1 in absolute value (with 0.09% were greater than 10).

### Attributions At Each t-SNE Iteration

For each experiment, we used the gradient of the t-SNE computed at step 250 for our attribution value (at the end of the early exaggeration phase). When we compared class-averaged attributions for each dataset, we found that attributions computed using gradients at this step best captured the globally-relevant features most often. This was especially apparent in our simulated data experiments, where the ground truth features were known. We hypothesize that this occurs due to the increased weight on the attractive forces during the early exaggeration phase. In the subsequent optimization steps, we found that the class-averaged gradients became uniform. We hypothesize that this is due to the increased weight on the repulsive forces. When this occurs the optimization adds more priority towards placing embeddings away from nearby neighboring ones. Inspecting the attributions computed during these later optimization steps may yield useful insights, but we leave such analysis for future work. See figure 7 for a visualization of the class-averaged attributions computed over each step of the t-SNE optimization.

**Fig. 7.**
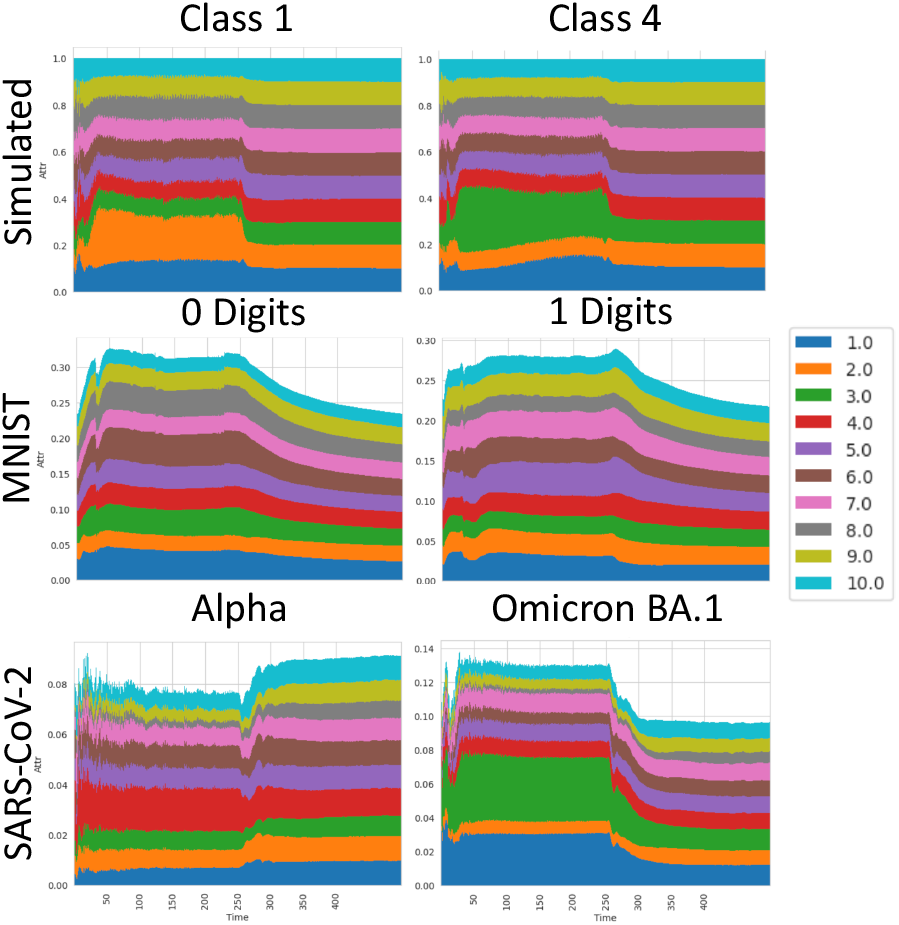
t-SNE Attributions at each step for each dataset. For each dataset, we select 2 classes displayed the (absolute value of the) class-averaged attributions across the first 500 steps of the t-SNE algorithm. We display the first 10 features (for datasets with more then 10 features, we ignore the rest here). (Top) The simulated dataset. We display Class 1 and 4. Note that for Class 1, attributions of feature 1 and 2 are significantly larger then the rest, as expected. Likewise for Class 4, the features with the largest attributions are the ground truth features. (Middle) For the 0 and 1 digit classes of MNIST, we see that certain features grow relative to the rest until step 250, after which the feature attribution sizes become increasingly uniform. (Bottom) We observe the same pattern for the SARS-CoV-2 data.

### Benchmarking Experiments

To assess the computational speed of our algorithm, we performed a series of benchmarking experiments on our t-SNE gradients implementation as well as our implementation of ordinary t-SNE. We performed this benchmarking experiment on synthetic datasets containing the same structure as described in figure 4. For all experiments, we only ran t-SNE for the first 250 steps, using the Barnes-Hut approximation. Both experiments were performed on a compute node provided by the Digital Research Alliance of Canada containing 16 CPU cores with 24GB RAM.

In our first experiment, we varied the number of features from 20 to 380 in increments of 20, keeping the number of samples fixed at 1000. In our second experiment, we varied the number of samples from 1000 to 20000 in increments of 1000, keeping the number of features fixed at 10. Not surprisingly, we found that both implementations scaled roughly loglinearly with respect to number of samples. As expected, we found that our gradient computations scaled roughly linearly with respect to number of input features. Note that the usual t-SNE is unaffected by the number of input features (except for the computation of *P* at the beginning), and so we did not benchmark this relationship. See figure 8 for more details.

**Fig. 8.**
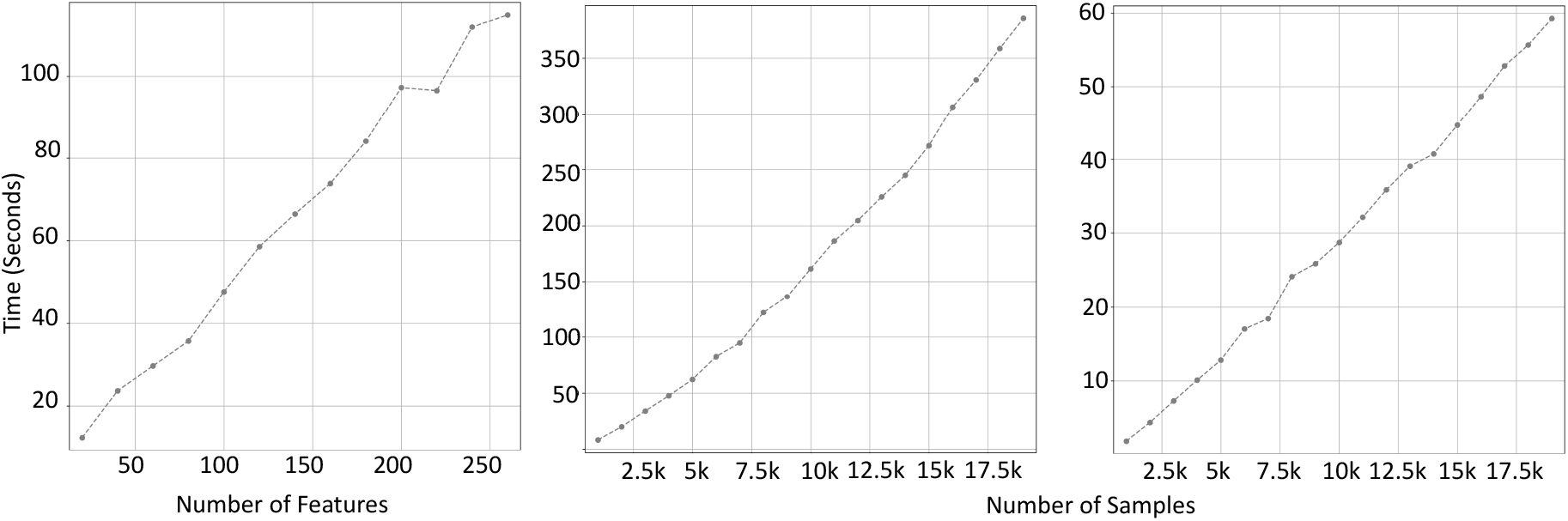
Benchmark experiments. (*Left*) complexity of our t-SNE attribution implementation with respect to the number of features. (*Center*) complexity with respect to the number of input samples. (*Right*) complexity with respect to number of input samples for t-SNE.

Qualitatively, we find that the attribution computation is considerably slower versus usual t-SNE. For example, our t-SNE implementation took around 2 seconds to compute, but the attributions took anywhere from 5 seconds to 2 minutes to compute. Despite this, our algorithm can in theory be applied to datasets of arbitrary size. This is since t-SNE is usually fitted on PC-transformed data with a small dimension of fixed size (usually 50-100). Therefore, the linear complexity with respect to the number of features should not be very noticeable in practice.

### Limitations and Unexpected Results of Gradient Attributions

We briefly discuss some limitations of our our t-SNE attribution method that we observed during our simulated data experiments. In the following two cases, the attributions did not identify the relevent features consistently (or at all):

1. When the dataset was small (*<* 1000 points).
2. When the feature enrichment was very large. For example the attributions of the simulated data with “Effect 1” = 8 performed poorly.

Furthermore, we note that, unexpectedly, the size of attributions for feature 1 was generally smaller then for feature 2 or 3, despite that “Effect 1” was always larger then “Effect 2”.

The cause of any of these limitations/unexpected results are not explored at depth here. We leave as future work a more rigorous characterization of these limitations and how to improve our method to resolve them.

## Competing interests

No competing interest is declared.

## Author contributions statement

MS developed the methodology and performed all analyses. JCG and RP pre-processed the SARS-CoV-2 data. MS wrote the paper, revised by JCG, RP, JGH. JGH and SL help interpret the results and supervised the project

## Acknowledgments

We would like to thank all laboratories that contributed to GISAID sequences. JGH is a Fonds de la Recherche du Québec en Santé (FRQS) Junior 2 Scholar. This project was funded by grants from NSERC (RGPIN-2022-04262) and the Institute for Data Valorization (CVD19-030) to JH. This study was also supported by the CIHR operating grant to the Coronavirus Variants Rapid Response Network (CoVaRR-Net).

## Supplemental Tables

We present the tables for the Synthetic data and MNIST attribution validation experiments here.

**Table S1.**
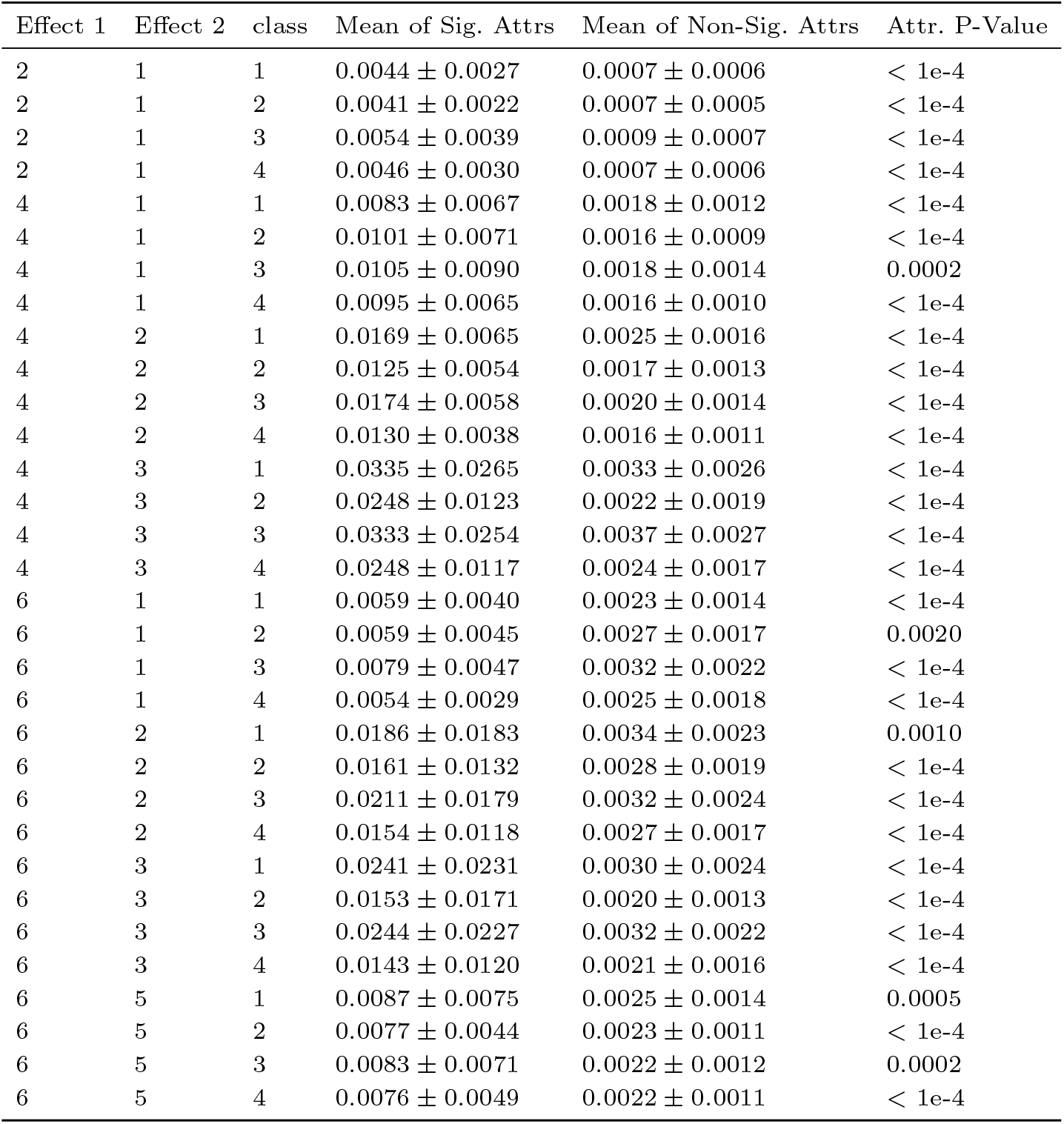
Results of synthetic data experiment

| Effect 1 | Effect 2 | class | Mean of Sig. Attrs | Mean of Non-Sig. Attrs | Attr. P-Value |
| --- | --- | --- | --- | --- | --- |
| 2 | 1 | 1 | $0.0044 \pm 0.0027$ | $0.0007 \pm 0.0006$ | $< 1e-4$ |
| 2 | 1 | 2 | $0.0041 \pm 0.0022$ | $0.0007 \pm 0.0005$ | $< 1e-4$ |
| 2 | 1 | 3 | $0.0054 \pm 0.0039$ | $0.0009 \pm 0.0007$ | $< 1e-4$ |
| 2 | 1 | 4 | $0.0046 \pm 0.0030$ | $0.0007 \pm 0.0006$ | $< 1e-4$ |
| 4 | 1 | 1 | $0.0083 \pm 0.0067$ | $0.0018 \pm 0.0012$ | $< 1e-4$ |
| 4 | 1 | 2 | $0.0101 \pm 0.0071$ | $0.0016 \pm 0.0009$ | $< 1e-4$ |
| 4 | 1 | 3 | $0.0105 \pm 0.0090$ | $0.0018 \pm 0.0014$ | 0.0002 |
| 4 | 1 | 4 | $0.0095 \pm 0.0065$ | $0.0016 \pm 0.0010$ | $< 1e-4$ |
| 4 | 2 | 1 | $0.0169 \pm 0.0065$ | $0.0025 \pm 0.0016$ | $< 1e-4$ |
| 4 | 2 | 2 | $0.0125 \pm 0.0054$ | $0.0017 \pm 0.0013$ | $< 1e-4$ |
| 4 | 2 | 3 | $0.0174 \pm 0.0058$ | $0.0020 \pm 0.0014$ | $< 1e-4$ |
| 4 | 2 | 4 | $0.0130 \pm 0.0038$ | $0.0016 \pm 0.0011$ | $< 1e-4$ |
| 4 | 3 | 1 | $0.0335 \pm 0.0265$ | $0.0033 \pm 0.0026$ | $< 1e-4$ |
| 4 | 3 | 2 | $0.0248 \pm 0.0123$ | $0.0022 \pm 0.0019$ | $< 1e-4$ |
| 4 | 3 | 3 | $0.0333 \pm 0.0254$ | $0.0037 \pm 0.0027$ | $< 1e-4$ |
| 4 | 3 | 4 | $0.0248 \pm 0.0117$ | $0.0024 \pm 0.0017$ | $< 1e-4$ |
| 6 | 1 | 1 | $0.0059 \pm 0.0040$ | $0.0023 \pm 0.0014$ | $< 1e-4$ |
| 6 | 1 | 2 | $0.0059 \pm 0.0045$ | $0.0027 \pm 0.0017$ | 0.0020 |
| 6 | 1 | 3 | $0.0079 \pm 0.0047$ | $0.0032 \pm 0.0022$ | $< 1e-4$ |
| 6 | 1 | 4 | $0.0054 \pm 0.0029$ | $0.0025 \pm 0.0018$ | $< 1e-4$ |
| 6 | 2 | 1 | $0.0186 \pm 0.0183$ | $0.0034 \pm 0.0023$ | 0.0010 |
| 6 | 2 | 2 | $0.0161 \pm 0.0132$ | $0.0028 \pm 0.0019$ | $< 1e-4$ |
| 6 | 2 | 3 | $0.0211 \pm 0.0179$ | $0.0032 \pm 0.0024$ | $< 1e-4$ |
| 6 | 2 | 4 | $0.0154 \pm 0.0118$ | $0.0027 \pm 0.0017$ | $< 1e-4$ |
| 6 | 3 | 1 | $0.0241 \pm 0.0231$ | $0.0030 \pm 0.0024$ | $< 1e-4$ |
| 6 | 3 | 2 | $0.0153 \pm 0.0171$ | $0.0020 \pm 0.0013$ | $< 1e-4$ |
| 6 | 3 | 3 | $0.0244 \pm 0.0227$ | $0.0032 \pm 0.0022$ | $< 1e-4$ |
| 6 | 3 | 4 | $0.0143 \pm 0.0120$ | $0.0021 \pm 0.0016$ | $< 1e-4$ |
| 6 | 5 | 1 | $0.0087 \pm 0.0075$ | $0.0025 \pm 0.0014$ | 0.0005 |
| 6 | 5 | 2 | $0.0077 \pm 0.0044$ | $0.0023 \pm 0.0011$ | $< 1e-4$ |
| 6 | 5 | 3 | $0.0083 \pm 0.0071$ | $0.0022 \pm 0.0012$ | 0.0002 |
| 6 | 5 | 4 | $0.0076 \pm 0.0049$ | $0.0022 \pm 0.0011$ | $< 1e-4$ |

**Table S2.** MNIST Experiment individual-level using mean averaging feature corruption

| Index | Correlation | Adjusted RAND Index | KNN Preservation |
| --- | --- | --- | --- |
| Random | $0.70 \pm 0.0715$ | $0.75 \pm 0.0290$ | $0.42 \pm 0.0017$ |
| Attribution $> 0$ | $0.58 \pm 0.0772$ | $0.62 \pm 0.0154$ | $0.33 \pm 0.0032$ |
| Attribution | $0.55 \pm 0.0587$ | $0.59 \pm 0.0261$ | $0.28 \pm 0.0035$ |
| Feature Value | $0.32 \pm 0.0510$ | $0.38 \pm 0.0129$ | $0.30 \pm 0.0006$ |
| Attribution · Feature Value | $0.35 \pm 0.0361$ | $0.36 \pm 0.0240$ | $0.20 \pm 0.0024$ |

**Table S3.**
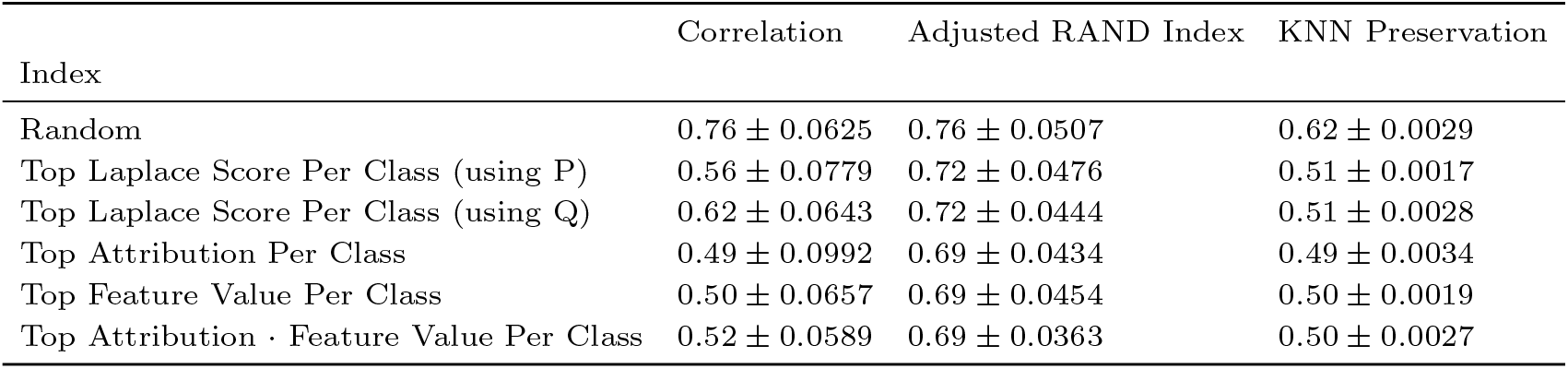
MNIST Experiment class-level using mean averaging feature corruption

**Table S4.** MNIST Experiment global-level using feature removal corruption

| Index | Correlation | Adjusted RAND Index | KNN Preservation |
| --- | --- | --- | --- |
| Random | $0.86 \pm 0.0326$ | $0.81 \pm 0.0458$ | $0.64 \pm 0.0037$ |
| Top Fisher Score | $0.68 \pm 0.0588$ | $0.72 \pm 0.0284$ | $0.54 \pm 0.0026$ |
| Top PC | $0.52 \pm 0.0521$ | $0.66 \pm 0.0425$ | $0.50 \pm 0.0020$ |
| Top Laplace Score (Using P) | $0.53 \pm 0.0509$ | $0.68 \pm 0.0408$ | $0.51 \pm 0.0013$ |
| Top Laplace Score (Using Q) | $0.53 \pm 0.0542$ | $0.68 \pm 0.0387$ | $0.51 \pm 0.0013$ |
| Top Feature Value | $0.52 \pm 0.0521$ | $0.66 \pm 0.0425$ | $0.50 \pm 0.0020$ |
| Attribution | $0.55 \pm 0.0665$ | $0.67 \pm 0.0434$ | $0.50 \pm 0.0042$ |
| Attribution · Feature Value | $0.51 \pm 0.0581$ | $0.67 \pm 0.0427$ | $0.50 \pm 0.0026$ |

**Table S5.** MNIST Experiment individual-level using permutation feature corruption

| Index | Correlation | Adjusted RAND Index | KNN Preservation |
| --- | --- | --- | --- |
| Random | $0.59 \pm 0.0760$ | $0.64 \pm 0.0189$ | $0.33 \pm 0.0011$ |
| Attribution $> 0$ | $0.46 \pm 0.0535$ | $0.51 \pm 0.0203$ | $0.23 \pm 0.0044$ |
| Attribution | $0.42 \pm 0.0457$ | $0.45 \pm 0.0320$ | $0.18 \pm 0.0043$ |
| Feature Value | $0.07 \pm 0.0140$ | $0.08 \pm 0.0026$ | $0.08 \pm 0.0004$ |
| Attribution · Feature Value | $0.13 \pm 0.0249$ | $0.14 \pm 0.0062$ | $0.08 \pm 0.0006$ |

**Table S6.** MNIST Experiment class-level using permutation feature corruption

| Index | Correlation | Adjusted RAND Index | KNN Preservation |
| --- | --- | --- | --- |
| Random | $0.75 \pm 0.0310$ | $0.77 \pm 0.0508$ | $0.47 \pm 0.0018$ |
| Top Laplace Score Per Class (using P) | $0.38 \pm 0.0263$ | $0.44 \pm 0.0350$ | $0.17 \pm 0.0009$ |
| Top Laplace Score Per Class (using Q) | $0.44 \pm 0.0313$ | $0.52 \pm 0.0413$ | $0.19 \pm 0.0022$ |
| Top Attribution Per Class | $0.28 \pm 0.0528$ | $0.42 \pm 0.0346$ | $0.18 \pm 0.0070$ |
| Top Feature Value Per Class | $0.18 \pm 0.0115$ | $0.26 \pm 0.0195$ | $0.13 \pm 0.0010$ |
| Top Attribution · Feature Value Per Class | $0.17 \pm 0.0301$ | $0.26 \pm 0.0191$ | $0.14 \pm 0.0038$ |

**Table S7.** MNIST Experiment global-level using permutation feature corruption

| Index | Correlation | Adjusted RAND Index | KNN Preservation |
| --- | --- | --- | --- |
| Random | $0.78 \pm 0.0357$ | $0.78 \pm 0.0483$ | $0.46 \pm 0.0022$ |
| Top Fisher Score | $0.57 \pm 0.0411$ | $0.62 \pm 0.0364$ | $0.24 \pm 0.0016$ |
| Top PC | $0.19 \pm 0.0321$ | $0.30 \pm 0.0207$ | $0.14 \pm 0.0014$ |
| Top Laplace Score (Using P) | $0.19 \pm 0.0302$ | $0.31 \pm 0.0240$ | $0.15 \pm 0.0009$ |
| Top Laplace Score (Using Q) | $0.19 \pm 0.0302$ | $0.31 \pm 0.0240$ | $0.15 \pm 0.0011$ |
| Top Feature Value | $0.19 \pm 0.0321$ | $0.30 \pm 0.0207$ | $0.14 \pm 0.0014$ |
| Attribution | $0.27 \pm 0.0485$ | $0.35 \pm 0.0401$ | $0.16 \pm 0.0076$ |
| Attribution · Feature Value | $0.20 \pm 0.0293$ | $0.30 \pm 0.0232$ | $0.15 \pm 0.0014$ |

## Footnotes

1 We include dimensionality reduction techniques such as t-SNE into our definition of ML models

2 intuitively, the larger the attribution, the less you have to change the corresponding input to achieve a fixed change in output

3 Note: *σ*_*i*_ is also a function of the input data. We ignore this relation when computing these gradients.

## Notes

https://github.com/MattScicluna/interpretable_tsne

### Competing Interest Statement

The authors have declared no competing interest.

### Summary of Updates

In the discussion, we clarify that: * our synthetic data experiments do not necessarily accurately represent real-world datasets and that attribution methods may not be appropriate if the data exhibits complicated patterns; * our method can be applied to synthetic single-cell RNA data such as Splatter: since we would have ground truth in simulated datasets, one could assess the effectiveness of our method on this specific data modality. We included additional sections in the appendix: * discussing the details of the t-SNE attribution computation, which includes a subsection discussing the complexity of the algorithm. We performed a series of benchmarking experiments to assess how our method scales with respect to sample size and number of features. * describing t-SNE attributions fitted on the Twenty Newsgroups Dataset: this additional experiment shows that our approach works on an additional modality (natural text) for which evaluation of attributions was possible.

